# A post-ER degradation pathway that relies on protease-dependent internalization from the vacuolar membrane

**DOI:** 10.1101/778530

**Authors:** Leticia Lemus, Zrinka Matić, Veit Goder

## Abstract

Newly synthesized proteins of the secretory pathway are quality-controlled inside the endoplasmic reticulum (ER) and, if not properly folded, are retained. An exception are glycosylphosphatidylinositol-anchored proteins (GPI-APs) which can leave the ER even when misfolded and are routed to the vacuole/lysosome for degradation by largely unknown mechanisms linked to post-ER quality control. Using yeast as model organism, we show that Gas1*, an ER-exported misfolded GPI-AP, is diverted from the secretory pathway to endosomes for transport to the vacuole. However, Gas1* is not sorted into endosomal intraluminal vesicles but internalizes directly from the vacuolar membrane. There, the vacuolar protease Pep4, but not any other known vacuolar protease, is required for Gas1* internalization. Our data reveal novel and unexpected mechanisms for invaginations from the vacuolar membrane.

**Highlights:**

- ER-exited misfolded GPI-anchored proteins are routed to the vacuole via endosomes but do not internalize into intraluminal vesicles
- Internalization occurs directly from the vacuolar membrane into intravacuolar mobile structures
- Internalization from the vacuolar membrane depends on the proteolytic activity of the vacuolar protease Pep4

## Introduction

A subclass of cell surface proteins facing the extracellular space in eukaryotes is attached to the membrane solely by a specific C-terminal glycolipid, the glycosylphatidylinositol (GPI) anchor. GPI-anchored proteins (GPI-APs) have important structural, developmental and signaling functions (Saha et al., 2015). They are initially synthesized with a C-terminal transmembrane domain (TMD) which is exchanged for a GPI anchor shortly after protein translocation into the endoplasmic reticulum (ER) and membrane insertion (Mayor and Riezman, 2004). After protein attachment, the GPI anchor undergoes extensive modifications by a process called remodeling. Remodeled GPI anchors are capable of binding to transmembrane adaptor molecules, such as the family of p24 proteins, which then promote ER exit of GPI-APs (Muniz et al., 2000; Fujita et al., 2009; Manzano-Lopez et al., 2015; Fujita et al., 2011; Castillon et al., 2009).

The remodeled GPI-anchor constitutes a strong ER export signal. In case protein folding fails, it can override ER protein quality control and retention mechanisms, leading to the ER export even of misfolded GPI-APs (Goder and Melero, 2011; Sikorska et al., 2016). In mammalian cells, rapid ER export of misfolded GPI-APs is enhanced upon induction of ER stress (Satpute-Krishnan et al., 2014). It is known that misfolded GPI-APs are ultimately degraded inside the vacuole in yeast and inside lysosomes (the equivalent of the vacuole) in mammalian cells (Satpute-Krishnan et al., 2014; Ashok and Hegde, 2008; Zavodszky and Hegde, 2019; Sikorska et al., 2016; Hirayama et al., 2008). However, most of their trafficking itinerary and the cellular mechanisms leading to their degradation, collectively called post-ER quality control, are unknown.

The vacuole/lysosome is the major hydrolytic compartment in eukaryotes and is required for the degradation and recycling of a vast number of substrates, including misfolded proteins. A major vacuolar protease in yeast is the endopeptidase Pep4, which is related to mammalian Cathepsin D (Parzych and Klionsky, 2019). It not only has broad substrate specificity but is also cleaving and thereby activating several other proteases inside the vacuole. Yeast cells with Pep4 deleted have strongly reduced vacuolar hydrolytic activity and can be used to preserve intravacuolar structures (Parzych and Klionsky, 2019).

The GPI anchor consists of lipids and carbohydrates. After remodeling, the lipid moieties consist of long chain fatty acids or ceramide. Degradation of a GPI-AP in its entirety, including its membrane-embedded GPI anchor, requires complete exposure to the interior of the vacuole/lysosome, and therefore requires protein internalization at some stage, a process that depends on membrane invagination. The best characterized pathway that leads to the internalization of transmembrane proteins for vacuolar/lysosomal degradation is the multi vesicular body (MVB) pathway, which is part of the endocytic pathway. The endocytic pathway transports membrane proteins and soluble cargo from the cell surface to the vacuole/lysosome. It also, however, receives material from the secretory pathway with which it intersects via membrane trafficking between the Golgi and endosomes (Conibear and Stevens, 1998). Among other things, this intersection serves to divert misfolded proteins from the early secretory pathway to endosomes. For instance, a misfolded mutant version of the yeast cell surface transmembrane protein Wsc1, called Wsc1*, escapes ER quality control and is diverted from the Golgi to endosomes. Through internalization from the endosomal membrane into intraluminal vesicles (ILVs) it becomes a cargo of MVBs and is then targeted to the vacuole (Wang et al., 2011). Alternatively, some misfolded proteins travel to the cell surface before being transported to the vacuole/lysosome through the endocytic pathway. This is the case for the misfolded version of the human prion protein PrP, called PrP*, a well-studied GPI-AP (Zavodszky and Hegde, 2019). For cargo molecules to be sorted into ILVs, they need to have TMDs for recognition and a cytosolic tail that contains lysine residues for modification with ubiquitin (Piper and Katzmann, 2007; Reggiori et al., 2000; Reggiori and Pelham, 2002; Hettema et al., 2004). When modified with ubiquitin, the tails are recognized by proteins of the endosomal sorting complexes required for transport (ESCRT) family which promote cargo concentration, de-ubiquitination, membrane invagination and ultimately lead to internalization of cargo molecules into ILVs (Schöneberg et al., 2016). Since GPI-APs lack both, TMDs and cytosolic tails, they diverge from typical MVB cargoes and their internalization into ILVs would depend on adaptor molecules. Interestingly, Cos proteins, belonging to the family of tetraspanins, are conserved multispanning membrane proteins that have been shown to possess adaptor functions for sorting cargo, including GPI-APs, into MVBs in yeast (MacDonald et al., 2015). Cos proteins operate exclusively in the endocytic pathway and modulate the metabolically regulated internalization of (correctly folded) cargo from endosomes. It is unknown if misfolded GPI-APs can rely on this mechanism for internalization into ILVs, either when coming from the secretory pathway or from the plasma membrane.

Microautophagy is a general term for vacuolar/lysosomal membrane protrusions or invaginations for the direct uptake of cytosol, large protein assemblies, (parts of) organelles such as the ER, or resident membrane proteins (Müller et al., 2000; Schuck et al., 2014; Roberts et al., 2003; Kiššová et al., 2007; van Zutphen et al., 2014; Oku and Sakai, 2018; Zhu et al., 2017). Invaginations from the vacuolar/lysosomal membrane were recently shown to require some ESCRT components and might thus be mechanistically related to the endosomal MVB pathway (Oku et al., 2017; Oku and Sakai, 2018). The extent to which ESCRT components are generally involved in membrane invaginations from the vacuole/lysosome is still unclear at present. Studies with yeast have shown the importance of additional factors, such as the VTC complex and the EGO complex, for various types of microautophagy (Uttenweiler et al., 2007; Dubouloz et al., 2005). Whether and how misfolded GPI-APs could exploit existing cellular pathways for the internalization directly from the vacuolar/lysosomal membrane for subsequent degradation is not known.

In summary, it is unclear how misfolded GPI-APs are routed to the vacuole/lysosome and what are the cellular mechanisms linked to their transport into the interior of the organelle. To address these issues, we have used the yeast *Saccharomyces cerevisiae* and have investigated the trafficking route and the degradation of the misfolded GPI-anchored protein Gas1*. With respect to wild type Gas1, which is a plasma membrane-localized glucanosyltransferase, Gas1* contains a single point mutation (G291R) which renders the protein misfolded, functionally impaired and subject to degradation inside the vacuole (Fujita et al., 2006; Sikorska et al., 2016).

## Results

### Misfolded GPI-APs enrich at the limiting membrane of the vacuole in Δpep4 cells

In order to follow the intracellular trafficking of Gas1* *in vivo*, we made use of a previously generated fusion protein that contains green fluorescent protein (GFP) downstream of the N-terminal signal sequence (Figure 1A and (Sikorska et al., 2016)). The construct, named GFP-Gas1*GPI, was integrated into the yeast genome and expressed under the control of the endogenous GAS1 promoter. When yeast lysate from wild type cells was analyzed by Western Blotting against GFP, full length GFP-Gas1*GPI as well as free GFP were detectable (Figures 1B, lane 1, and 1C). GFP fusion proteins generate free GPF inside the vacuole, due to the stable fold of the GFP moiety that leads to a retardation of its degradation (Guimaraes et al., 2015). The mutant *Δpep4*, with strongly reduced vacuolar hydrolytic activity, produced only negligible amounts of free GFP (Figures 1B, lane 2, and 1C). Together these results confirm that GFP-Gas1*GPI is routed to the vacuole, as reported previously (Sikorska et al., 2016). We then analyzed the trafficking of GFP-Gas1*GPI by live cell fluorescence microscopy, in combination with differential interference contrast (DIC) imaging for the easy identification of the vacuole (Figure 1D). In wild type cells, most of the fluorescent signal was detected inside the vacuole with little or no labeling of the ER at steady state, in agreement with rapid ER exit of GFP-Gas1*GPI (Figure 1D, wild type). In *Δpep4* cells, labeling of the perinuclear ER became stronger compared to wild type cells, suggesting an increased ER retention of the protein if vacuole function is compromised (Figure 1D, *Δpep4*, “*pER*”). Most interestingly, however, we detected a striking reduction of fluorescence in the vacuolar lumen in *Δpep4* cells compared to wild type cells and observed an enrichment of fluorescence in the limiting vacuolar membrane (Figure 1D, *Δpep4*, arrow heads). Laser scanning confocal fluorescence microscopy confirmed these results (Figure 1E). In order to test whether this is a more general phenomenon we generated a distinct misfolded GPI-AP, named GFP-CPY*GPI (Figure S1A). Again, this protein accumulated in the vacuolar membrane in *Δpep4* cells (Figure S1B).

**Figure 1.**
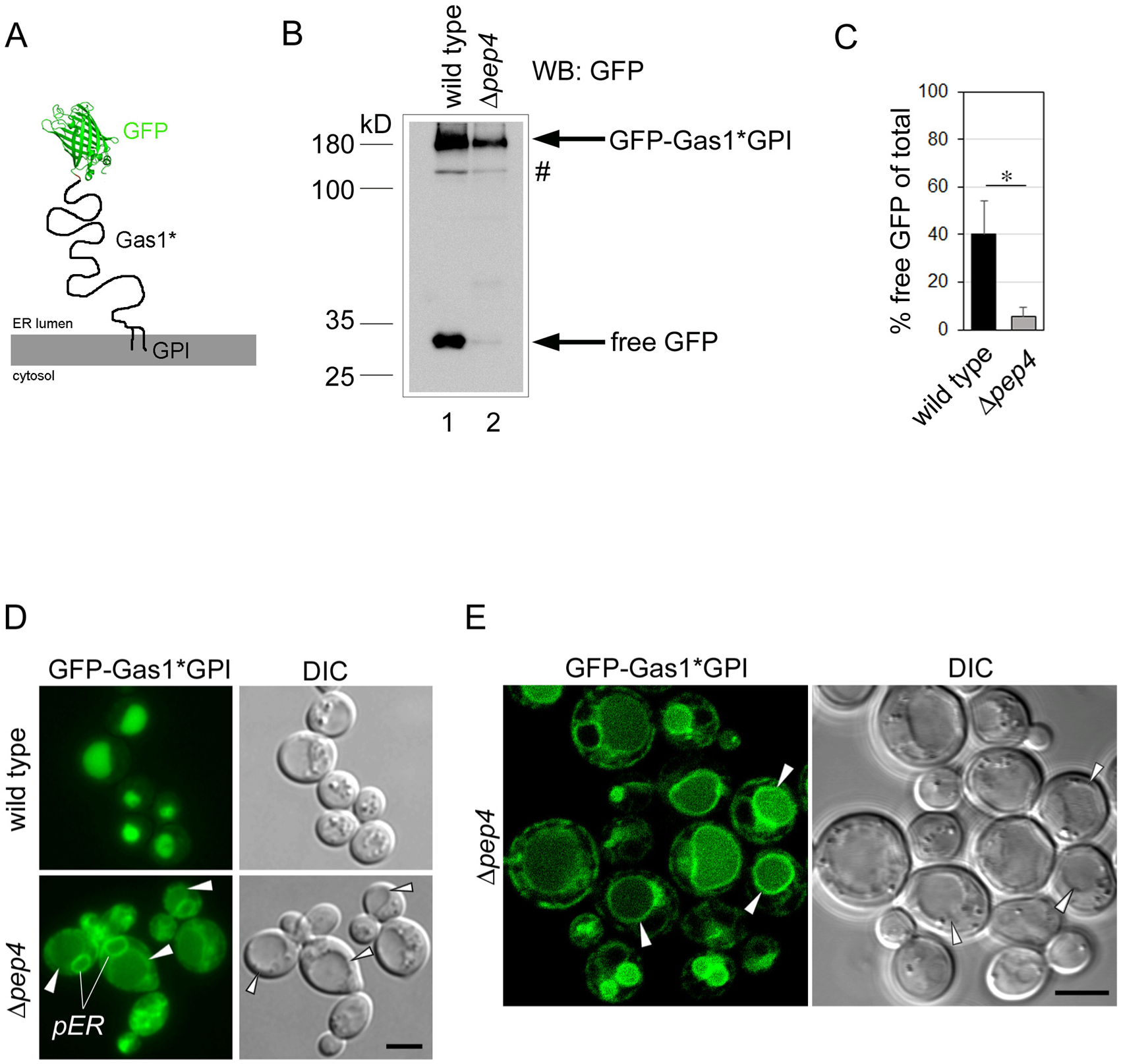
Misfolded GPI-APs enrich at the limiting membrane of the vacuole in *Δpep4* cells. **(A)** Schematic representation of the construct GFP-Gas1*GPI. The GFP moiety was fused to the N-terminal domain of Gas1*, downstream of the signal sequence, and faces the ER lumen. The C-terminal GPI anchor extends into the luminal leaflet of the ER membrane. **(B)** GFP cleavage assay. Wild type cells and *Δpep4* cells expressing genomically integrated GFP-Gas1*GPI under the control of the endogenous GAS1 promoter were lysed and analyzed by SDS-PAGE in combination with Western Blotting (WB) with antibodies against GFP. Hashtag represents a minor fraction of the fusion protein that likely has not been translocated into the ER and is therefore not N-glycosylated. **(C)** Quantification and statistical analysis of results from experiments shown in (B). Mean values and standard deviations from 3 individual experiments are shown. *p˂0.05 (unpaired two-tailed Student’s t-test). **(D)** Cells used for Western Blot analysis in (B) were analyzed by live cell fluorescence microscopy and differential interference contrast (DIC) microscopy. Arrow heads indicate the limiting membrane of the vacuole. *pER* = perinuclear ER. Scale bar: 5μm. **(E)** *Δpep4* cells were analyzed by live cell laser scanning confocal fluorescence microscopy and DIC microscopy. Arrow heads indicate the limiting membrane of the vacuole. Scale bar: 5μm. **See also Figure S1.**

These results combined suggest that misfolded GPI-APs are routed to the limiting membrane of the vacuole prior to degradation inside the vacuolar lumen.

### Generation and vacuolar uptake of ILVs and autophagosomes is functional in cells that express GFP-Gas1*GPI

The same results predict that neither the MVB pathway nor autophagy is routing GPI-APs into the vacuole, since both pathways are known to be fully functional in *Δpep4* cells (Xie et al., 2008; Gong and Chang, 2001; Wang et al., 2011). In support of this, GFP-Gas1*GPI targeting to the vacuole was not affected in mutant cells that are defective in generating autophagosomes (Figure S2).

Since GPI anchors alter membrane composition (Saha et al., 2015), we also considered the possibility that the constitutive expression of a misfolded GPI-AP might generally affect the formation of ILVs and/or autophagosomes and their uptake by the vacuole. In order to test this, we co-expressed GFP-Gas1*GPI together with substrates and markers of either the MVB pathway or autophagy in *Δpep4* cells. We first co-expressed GFP-Gas1*GPI together with mRFP-Ufe1 (Figure 2A). We have previously shown that the <u>s</u>oluble <u>N</u>SF <u>a</u>ttachment protein <u>re</u>ceptor (SNARE) Ufe1, a normally ER-localized membrane protein, becomes a substrate of the MVB pathway when overexpressed or if autophagy is induced (Lemus et al., 2016). In wild type cells, both proteins are routed into the vacuolar lumen, as expected (Figure 2B, wild type). In *Δpep4* cells, mRFP-Ufe1 is still targeted to the lumen of the vacuole, with no accumulation in the vacuolar membrane detectable, whereas GFP-Gas1*GPI was found enriched in the vacuolar membrane (Figure 2B, *Δpep4*).

**Figure 2.**
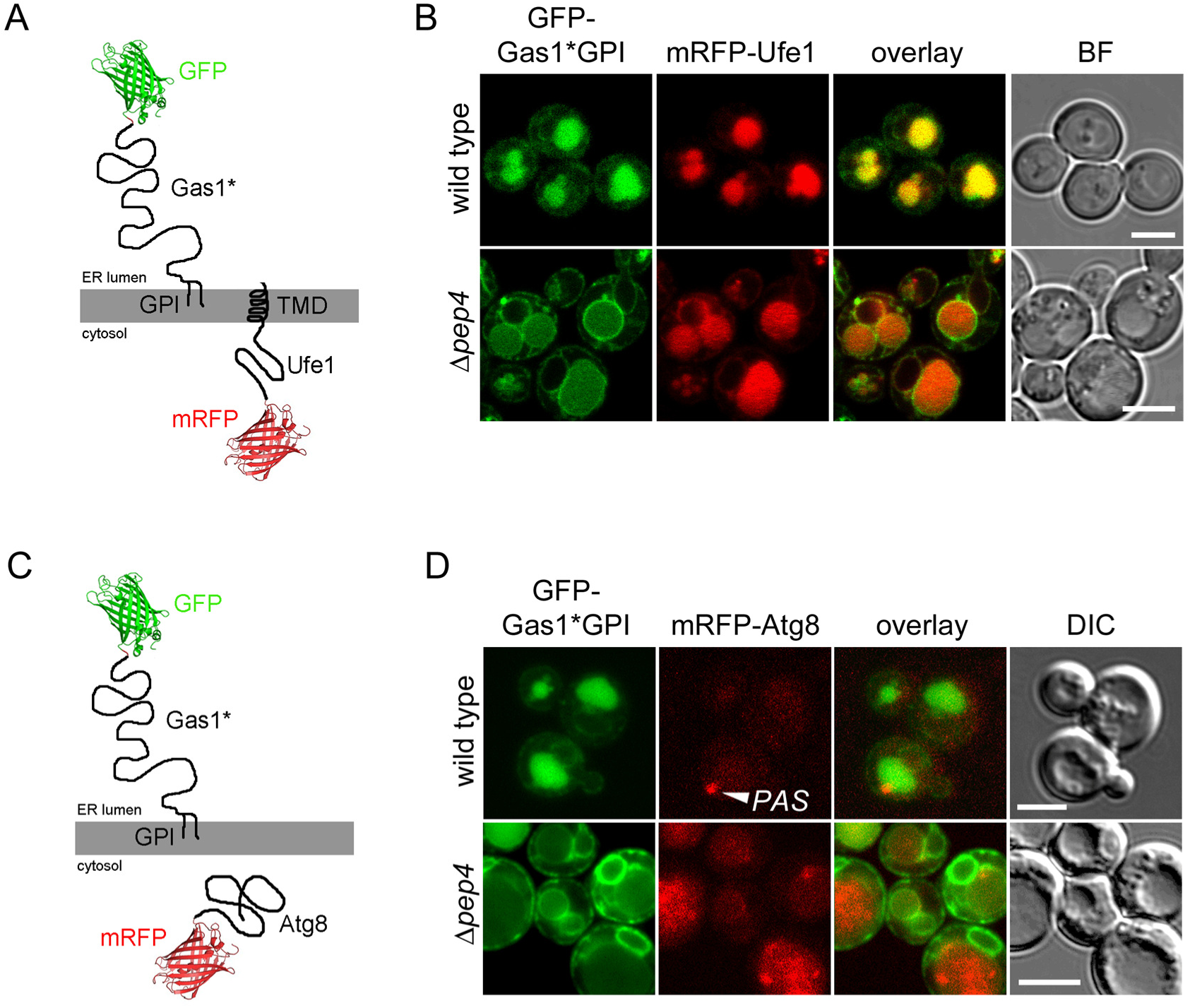
Generation and vacuolar uptake of ILVs and autophagosomes is functional in cells that express GFP-Gas1*GPI. **(A)** Schematic representation of the constructs used in the experiment shown in (B). GFP-Gas1*GPI and mRFP-Ufe1 have the opposite orientation in the ER membrane as indicated. Ufe1 possesses a C-terminal transmembrane domain (TMD). **(B)** Wild type cells and *Δpep4* cells co-expressing genomically integrated GFP-Gas1*GPI under the control of the GAS1 promoter and mRFP-Ufe1 expressed from a CEN plasmid under the control of the TDH3 promoter were analyzed by live cell laser scanning confocal fluorescence microscopy. BF=bright field image. Scale bar: 5μm. **(C)** Schematic representation of the constructs used in the experiment shown in (D). GFP-Gas1*GPI and mRFP-Atg8 have distinct cellular localizations as indicated. **(D)** Wild type cells and *Δpep4* cells co-expressing genomically integrated GFP-Gas1*GPI under the control of the GAS1 promoter and mRFP-Atg8 expressed from a CEN plasmid under the control of its endogenous promoter were analyzed like in (B). *PAS* = pre-autophagosomal structure. Scale bar: 5μm.

Next, we co-expressed GFP-Gas1*GPI together with the autophagosomal marker mRFP-Atg8, a soluble cytosolic protein (Figure 2C). Under nutrient-rich conditions, Atg8 is constitutively incorporated into <u>c</u>ytoplasm to <u>v</u>acuole targeting (Cvt) vesicles with a diameter between 140 and 160 nm. Cvt vesicles resemble selective autophagosomes for the transport of specific vacuolar proteases from the cytosol to the vacuole, where they are rapidly dissolved (Lynch-Day and Klionsky, 2010). mRFP-Atg8 is occasionally visible as a perivacuolar dot-like structure, called the PAS, which resembles the assembly site for Cvt vesicles in the cytosol (Figure 2D, “PAS”). In wild type cells, both proteins are expected to be routed to the vacuole interior (Figure 2D, wild type). However, only GFP-Gas1*GPI was clearly visible inside the vacuole, whereas the low expression level of mRFP-Atg8 under growing conditions only produced a diffuse and very faint signal across the entire cell, except for the PAS (Figure 2D, wild type). Importantly, however, in *Δpep4* cells, where Cvt vesicles are known to be structurally preserved inside the vacuole (Baba et al., 1997), we could clearly detect dot-like mRFP-Atg8-containing structures inside the vacuole, with no accumulation in the vacuolar membrane (Figure 2D, *Δpep4*). In contrast, GFP-Gas1*GPI was found enriched in the vacuolar membrane (Figure 2D, *Δpep4*).

These results combined show that expression of a misfolded GPI-AP does not inhibit the generation and transport of ILVs and autophagosomes in *Δpep4* cells. In conclusion, GFP-Gas1*GPI is transported to the interior of the vacuole through an uncharacterized pathway that is independent of MVBs and autophagosomes.

### Misfolded Gas1* but not correctly folded Gas1 is diverted from the secretory pathway to endosomes and routed to the vacuole in an ESCRT-dependent manner

Since our reporter is routed into the vacuole by a mechanism that differs from that for an MVB substrate, it could indicate that misfolded GPI-APs bypass endosomes altogether. Known trafficking routes from the ER to the vacuole are the CPY pathway and the ALP pathway (Raymond et al., 1992; Piper et al., 1995). The former routes substrates from the ER to the vacuole via the Golgi and endosomes, the latter is a pathway from the ER to the vacuole via only the Golgi, bypassing endosomes. When we tested whether routing of GFP-Gas1*GPI to the vacuole was affected in *Δapm3* cells, which specifically inhibit the ALP pathway (Cowles et al., 1997), we did not observe any intracellular accumulations of GFP-Gas*GPI when compared with wild type cells (Figure S3). We then tested whether routing of GFP-Gas1*GPI to the vacuole is affected in ESCRT mutants, which specifically inhibit the CPY pathway between endosomes and the vacuole (Katzmann et al., 2001, 2003; Babst et al., 2002). In all ESCRT mutants, GFP-Gas1*GPI accumulated in enlarged perivacuolar structures, known as the prevacuolar compartment (PVC), and was not visible inside the vacuole (Figure 3A, GFP-Gas1*GPI). In addition, all mutants showed a reduction in the generation of free GFP (Figures 3B and C). The accumulation of the reporter in the PVC is a typical phenotype for substrates that are routed through the CPY pathway and supports the notion that GFP-Gas1*GPI is transported to the vacuole via endosomes. Interestingly, we did not observe labeling of the vacuolar membrane with our reporter in the tested ESCRT mutants.

**Figure 3.**
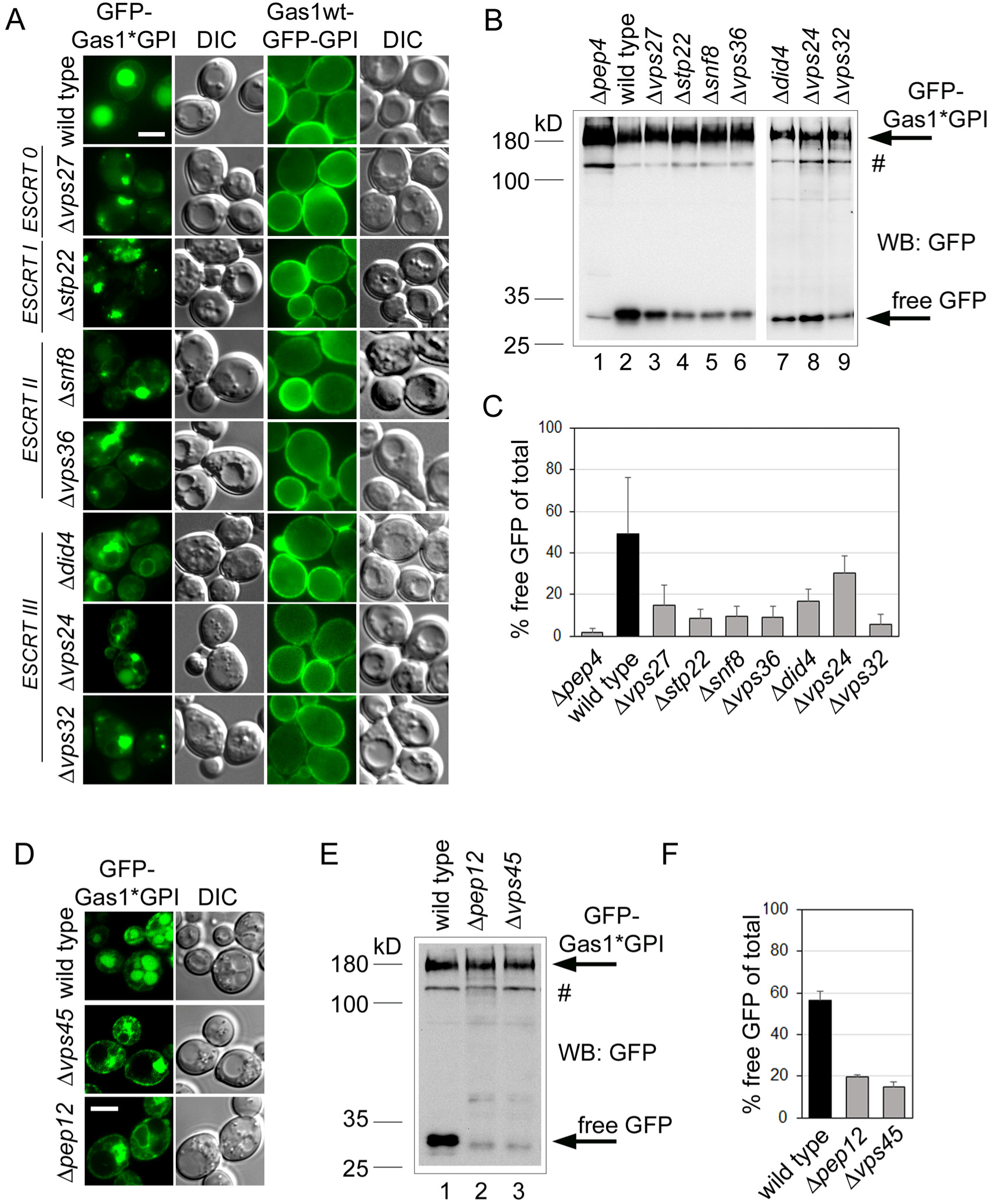
Misfolded Gas1* but not correctly folded Gas1 is diverted from the secretory pathway to endosomes and routed to the vacuole in an ESCRT-dependent manner. **(A)** Wild type cells and the indicated mutant cells expressing misfolded GFP-Gas1*GPI or correctly folded Gas1wt-GFP-GPI from a CEN plasmid under the control of the GAS1 promoter were analyzed by live cell fluorescence microscopy and DIC microscopy. The mutants belong to the indicated classes of ESCRT genes. Scale bar: 5μm. **(B)** GFP cleavage assay with wild type cells and the mutant cells used in (A) expressing GFP-Gas1*GPI. Cells were lysed and analyzed by SDS-PAGE in combination with Western Blotting (WB) with antibodies against GFP. Hashtag represents a minor fraction of the fusion protein that likely has not been translocated into the ER. **(C)** Quantification and statistical analysis of results from experiments shown in (B). Mean values and standard deviations from 3 individual experiments are shown. **(D)** Wild type cells and the indicated mutant cells expressing misfolded GFP-Gas1*GPI were analyzed by live cell fluorescence microscopy and DIC microscopy. Scale bar: 5μm. **(E)** GFP cleavage assay with wild type cells and the mutant cells used in (D). **(F)** Quantification and statistical analysis of results from experiments shown in (D). Mean values and standard deviations from 3 individual experiments are shown. **See also Figure S3.**

In a control experiment we used GFP-tagged wild type Gas1, where GFP was fused with Gas1 just upstream of the GPI anchor attachment site. In wild type cells, Gas1wt-GFP-GPI is trafficked to the cell surface, as expected (Figure 3A, Gas1wt-GFP-GPI, wild type).We observed no alteration in plasma membrane labeling in the ESCRT mutants when compared with wild type cells, showing that the wild type protein does not require the ESCRT machinery for trafficking and is not routed through endosomes (Figure 3A, Gas1wt-GFP-GPI).

We then tested whether the transport of GFP-Gas1*GPI to endosomes occurs directly from the Golgi or indirectly, via initial targeting from the Golgi to the plasma membrane followed by endocytosis, like observed for PrP* in mammalian cells (Satpute-Krishnan et al., 2014; Zavodszky and Hegde, 2019). We did occasionally observe plasma membrane labeling with GFP-Gas1*GPI in wild type cells at steady state, albeit very faint. When we used *Δvps45* and *Δpep12* cells, which specifically inhibit direct transport from the Golgi to endosomes but leave transport to endosomes via the plasma membrane intact (Bryant et al., 1998), we observed that trafficking to the vacuole was almost completely blocked (Figure 3D). In addition, the generation of free GPF was strongly reduced in these mutants (Figures 3E and F).

In summary, these results suggest that the majority of GFP-Gas1*GPI is routed directly from the Golgi to endosomes and, if any, only a minor fraction is routed to endosomes via the plasma membrane. GFP-Gas1*GPI is subsequently transported in endosomes to the vacuole and this transport step strictly depends on the ESCRT machinery.

### Pep4-dependent protein internalization from the vacuolar membrane

Despite the dependency on ESCRT proteins for its routing to the vacuole, GFP-Gas1*GPI does not internalize into ILVs. These results could suggest that misfolded proteins with exclusively luminal exposure and lacking TMDs, such as GPI-APs, have no need to internalize and are degraded directly from the vacuolar membrane by Pep4 or by vacuolar proteases that depend on Pep4 activity. However, given that the membrane-embedded remodeled GPI anchor contains long chain fatty acids or ceramide, it is likely that additional mechanisms such as invaginations or membrane recycling must exist in order to maintain vacuolar membrane lipid homeostasis. Intravacuolar membrane-bound structures like microautophagic vesicles, Cvt vesicles, autophagic bodies and ILVs are rapidly dissolved by the action of Atg15, a vacuolar lipase (Teter et al., 2001; Ramya and Rajasekharan, 2016). As such, an intravacuolar vesicle that originates from invagination from the vacuolar membrane is likely also sensitive to Atg15. We therefore expressed GFP-Gas1*GPI in *Δatg15* and *Δpep4Δatg15* mutant cells but could not detect intravacuolar fluorescent structures (not shown).

Given the topology of GFP-Gas1*GPI, intravacuolar vesicles that contain the reporter would have GFP face the vacuolar lumen. Full vacuolar protease activity in *Δatg15* cells and residual protease activity in *Δpep4Δatg15* cells might lead to a rapid release of the fluorescent moiety from the membrane and thus prevent the detection of intravacuolar structures. We therefore generated a modified reporter. We deleted the N-terminal signal sequence of GFP-Gas1*GPI and introduced the signal anchor of the ER protein Ted1 in between GFP and the downstream coding sequence of Gas1*. The resulting chimeric protein, named GFP-TMD-Gas1*GPI, has its GFP moiety on the cytosolic side of the membrane but retains the luminal localization of Gas1* (Figure 4A). Internalization of this reporter from the vacuolar membrane would trap GFP inside vesicles and prevent its proteolytic release from intravacuolar vesicles until their membranes are dissolved.

**Figure 4.**
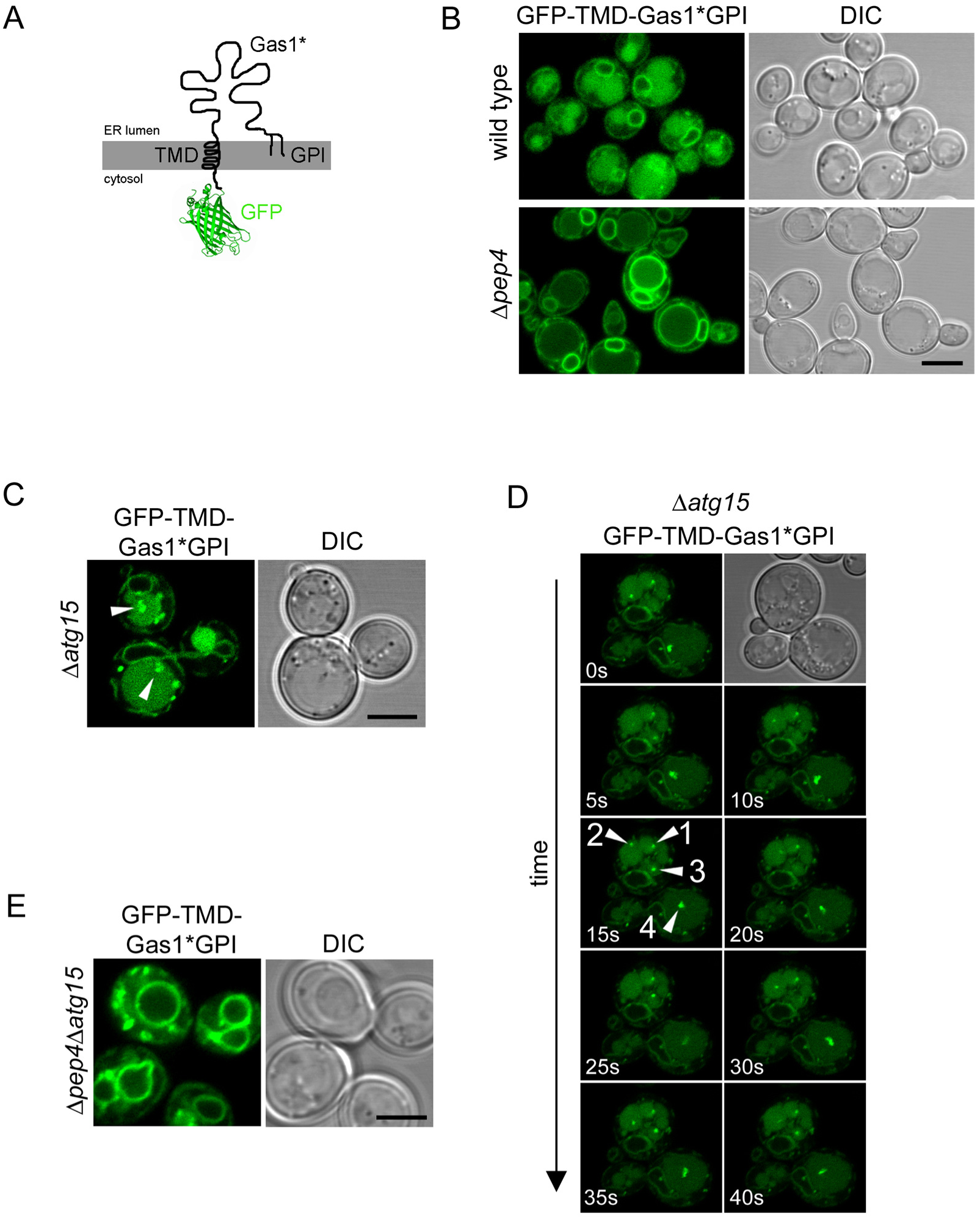
Pep4-dependent protein internalization from the vacuolar membrane. **(A)** Schematic representation of the construct GFP-TMD-Gas1*GPI. The GFP moiety is localized to the cytosol and connected via a TMD to luminal Gas1*GPI. **(B, C and E)** Wild type cells and the indicated mutant cells expressing GFP-TMD-Gas1*GPI from a CEN plasmid under the control of the GAS1 promoter were analyzed by live cell laser scanning confocal fluorescence microscopy and DIC microscopy. Arrow heads in (C) indicate intravacuolar fluorescent structures. Scale bar: 5μm. **(D)** Individual frames from time lapse imaging with *Δatg15* cells expressing GFP-TMD-Gas1*GPI. Numbers in lower left corners indicate the time in seconds after start of the image sequence. Arrow heads indicate 4 selected intravacuolar fluorescent structures. Structure 1 is immobile during the time lapse imaging, whereas structures 2, 3 and 4 are mobile. **See also Figure S4 and Movies S1 and S2.**

Expression of the reporter in wild type cells led to a significant amount of GFP being detectable inside the vacuolar lumen (Figure 4B, wild type). Compared with GFP-Gas1*GPI, the TMD-containing reporter shows more labeling of the perinuclear ER, indicative of increased ER retention, probably due to the presence of the TMD of the ER-resident protein Ted1 (Figure 4B, wild type; compare with Figure 1D). When GFP-TMD-Gas1*GPI was expressed in *Δpep4* cells, we observed a dramatic loss of labeling of the vacuolar lumen and detected labeling of the vacuolar membrane, similar to GFP-Gas1*GPI (Figure 4B, *Δpep4*; compare with Figure 1D). Thus, GFP-TMD-Gas1*GPI and GFP-Gas1*GPI share the same phenotype with respect to accumulation in the vacuolar membrane. When we expressed GFP-TMD-Gas1*GPI in *Δatg15* cells, we now detected fluorescent structures inside the vacuolar lumen (Figure 4C, arrows). These structures were mobile and, based on their distinct velocity, had different diameters (Figure 4D and Movie S1). Importantly, *Δatg15* cells did not show labeling of the vacuolar membrane, suggesting that the intravacuolar fluorescent structures result from the material that is seen in the vacuolar membrane in *Δpep4* cells (Figures 4C and D, compare with Figure 4B, *Δpep4*). Indeed, when we expressed GFP-TMD-Gas1*GPI in *Δpep4Δatg15* cells, we observed that the generation of intravacuolar fluorescent structures was completely suppressed and the vacuolar membrane was strongly labeled (Figure 4E and Movie S2). These results suggest that *Δpep4* cells have a deficiency in internalization of all our tested reporters from the vacuolar membrane.

The mechanism of internalization observed with GFP-TMD-Gas1*GPI could resemble known processes linked to microautophagy. We therefore tested whether previously identified components with functions in various types of microautophagy would affect the internalization of GFP-Gas1*GPI, such as components of the VTC complex and the EGO complex (Uttenweiler et al., 2007; Dubouloz et al., 2005). None of the tested deletion mutants showed an enrichment of GFP-Gas1*GPI in the limiting vacuolar membrane, all mutants showed labeling of the vacuolar lumen, like wild type cells (Figure S4A).

In an additional set of experiments, we used a construct where the C-terminal GPI anchor of GFP-TMD-Gas1*GPI was converted into a second TMD (Figure S4B). This construct, named GFP-TMD-Gas1*TMD, resulted in strong labeling of the perinuclear ER and strong reduction in routing of the reporter to the vacuole in wild type cells, supporting the idea that the ER exit of GFP-TMD-Gas1*GPI is driven by the GPI anchor, as expected (Figure S4C and (Sikorska et al., 2016)). Interestingly, a minor fraction of GFP-TMD-Gas1*TMD left the ER and was still accumulating in the vacuolar membrane in *Δpep4* cells (Figure S4C). This shows that the failure of protein internalization is not exclusively linked to misfolded proteins that contain a GPI anchor.

### From all known and predicted vacuolar proteases only Pep4 is required for internalization of GFP-Gas1*GPI from the vacuolar membrane

Pep4 is a general activator of vacuolar proteases and its deletion causes therefore a drop in proteolytic activity inside the vacuole (Parzych and Klionsky, 2019). In order to address whether the internalization defect of GFP-Gas1*GPI in *Δpep4* cells is linked to the activity of Pep4 alone or to a specific vacuolar protease that depends directly or indirectly on activation by Pep4 we tested the contribution of all known and predicted vacuolar proteases to the internalization of the reporter. Of the nine well-characterized vacuolar proteases, only the deletion of Pep4 resulted in accumulation of the reporter in the vacuolar membrane (Figure 5A). In addition, of five genes that are predicted to code for yet uncharacterized vacuolar proteases (Parzych and Klionsky, 2019; Yofe et al., 2016), none produced vacuolar membrane labeling with GFP-Gas1*GPI when deleted (Figure 5B). Vacuolar localization of the predicted proteases was verified with live cell fluorescence microscopy using mCherry fusion proteins (Figure S5). In summary, our data show that the internalization defect obtained with *Δpep4* cells is not linked to other known or predicted vacuolar proteases.

**Figure 5.**
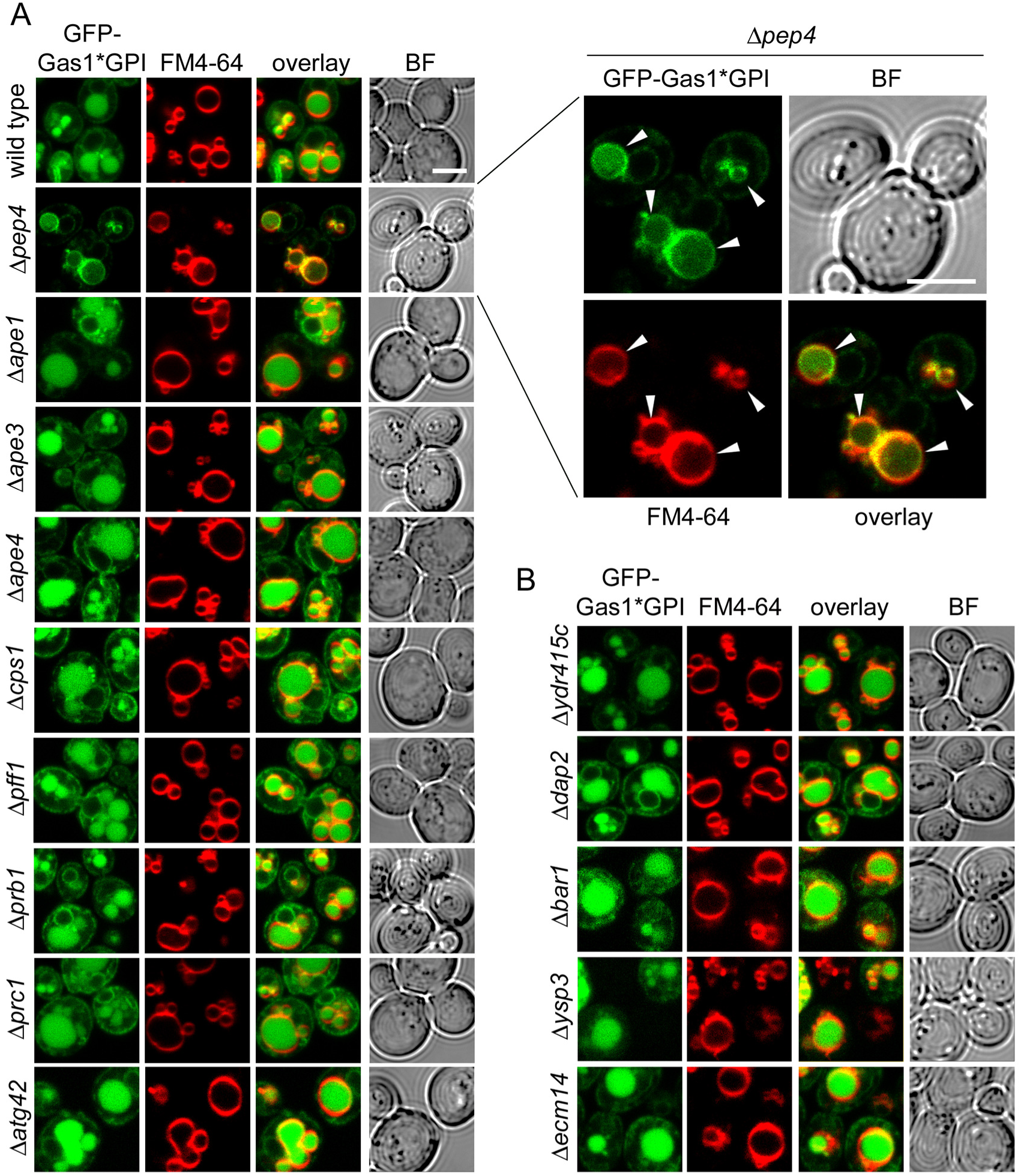
From all known and predicted vacuolar proteases only Pep4 is required for internalization of GFP-Gas1*GPI from the vacuolar membrane. **(A)** Wild type cells and the indicated deletion mutant cells expressing GFP-Gas1*GPI from a CEN plasmid under the control of the GAS1 promoter were analyzed by live cell fluorescence microscopy in combination with staining of the vacuolar membrane with FM4-64. The set of the presented images taken with *Δpep4* cells is additionally shown after enlargement. Arrow heads indicate the co-labeling of the vacuolar membrane with FM4-64 and GFP. Scale bar: 5μm. BF = bright field. **(B)** Same like in (A) with additional deletion mutants.

### The proteolytic activity of Pep4 is required for internalization of GFP-TMD-Gas1*GPI

These results prompted us to test directly whether the protease activity of Pep4 is required for reporter internalization. We used *Δpep4Δatg15* cells and transformed them with either an empty plasmid or with a plasmid that contained either myc-tagged wild type Pep4 or a catalytically inactive myc-tagged mutant (D109K) version of the protease, both expressed under the control of the endogenous promoter (Figure 6A). In the presence of wild type Pep4, but not in presence of either the empty plasmid or the catalytically inactive mutant, we could observe the disappearance of the characteristic labeling of vacuolar membrane and the appearance of intravacuolar vesicles containing GFP-TMD-Gas1*GPI (Figure 6A). Expression of both myc-tagged versions of Pep4 was verified by Western Blotting (Figure 6B). We noticed that Pep4(D109K)-myc ran with a slightly elevated molecular weight compared to the Pep4(wt)-myc and showed increased steady-state levels (Figure 6B, compare lanes 2 and 3). This is likely a result of reduced self-processing and reduced vacuolar degradation capacity (Parzych and Klionsky, 2019). In summary, our results show that the proteolytic activity of Pep4 is required for internalization of GFP-TMD-Gas1*GPI from the vacuolar membrane.

**Figure 6.**
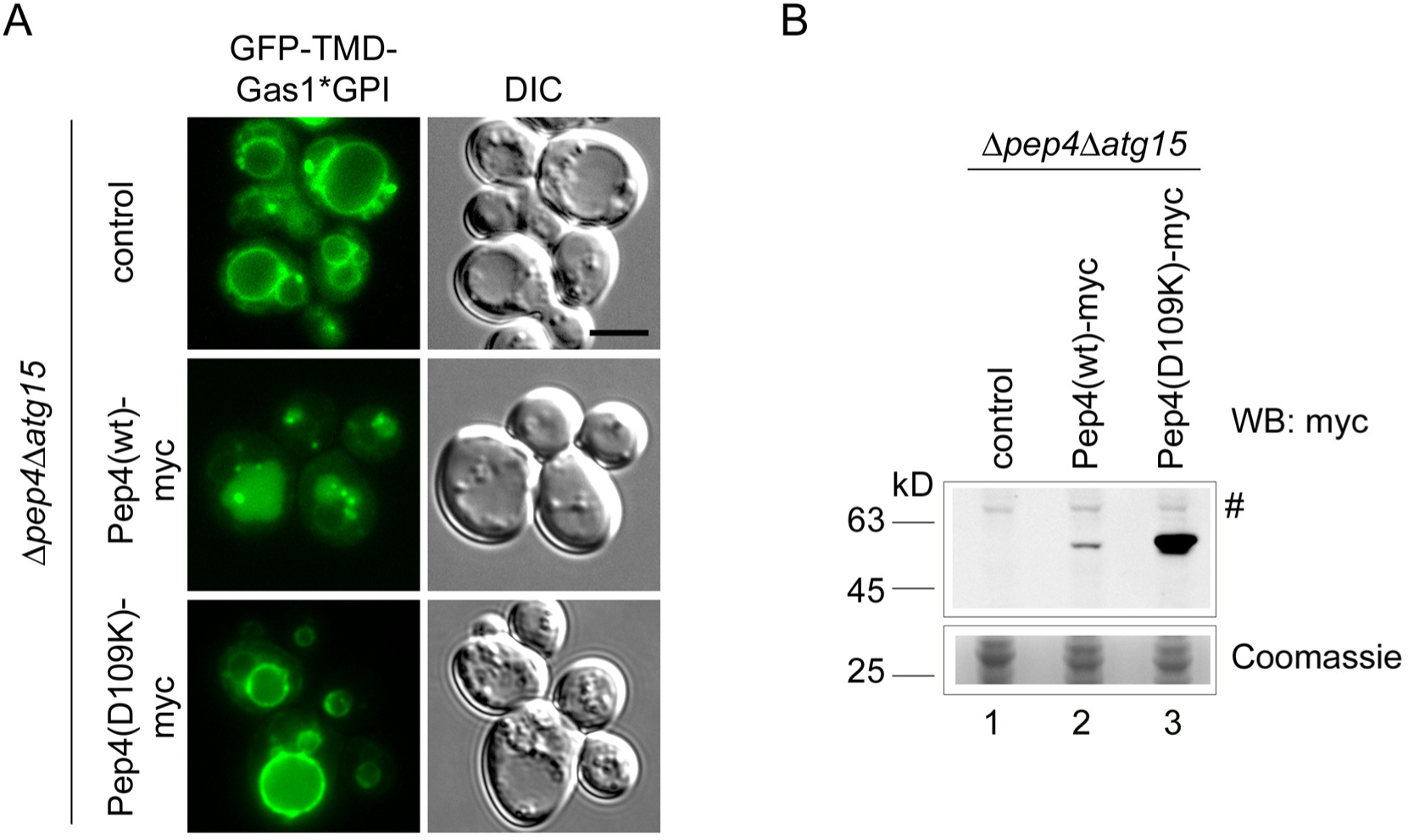
The proteolytic activity of Pep4 is required for internalization of GFP-TMD-Gas1*GPI. **(A)** *Δpep4Δatg15* cells expressing GFP-TMD-Gas1*GPI from a CEN plasmid under the control of the GAS1 promoter were transformed with a 2 micron plasmid containing either no insert (control), or myc-tagged Pep4, or catalytically inactive myc-tagged Pep4(D109K), both under the control of its endogenous promoter. Cells were analyzed by live cell laser scanning confocal fluorescence microscopy and DIC microscopy. Scale bar: 5μm. **(B)** Cells used in (A) were lysed and analyzed by SDS-PAGE in combination with Western Blotting (WB) with antibodies against the myc tag. Hashtag represents an unspecific band that is useful to indicate the slightly higher molecular weight of Pep4(D109K)-myc (lane 3) compared to Pep4(wt)-myc (lane 2).

## Discussion

By addressing the processes linked to the trafficking and degradation of misfolded GPI-APs we have discovered novel cellular mechanisms that regulate membrane traffic and vacuolar membrane dynamics.

First, our study delineates the post-ER degradation pathway of the misfolded GPI-AP Gas1* (Figure 7). We found that transport of Gas1* to the vacuole depends on Vps45 and Pep12, components that regulate protein sorting and trafficking from the Golgi to endosomes, also termed the CPY pathway (Raymond et al., 1992; Piper et al., 1995). Alternative pathways for the transport from the Golgi to the vacuole bypassing endosomes (AP-3 pathway) and routing to the plasma membrane followed by endocytosis play no or only a minor role. Thus, we propose that Gas1* is routed initially to the Golgi along the early secretory pathway from where it is diverted to endosomes. In contrast, wild type Gas1 bypasses endosomes on its way to the plasma membrane (Figure 7). These data are in agreement with previous reports about the relevance of the Golgi in post-ER quality control and in directing substrates to the vacuole for degradation (Arvan et al., 2002; Vashist et al., 2001; Wang and Ng, 2010; Watanabe et al., 2019; Dobzinski et al., 2015). It is unclear at present which signal is recognized inside the Golgi and what mechanism diverts Gas1* to endosomes. We found that subsequent trafficking of Gas1* in endosomes to the vacuole is strictly dependent on ESCRT components. We therefore propose that endosomes carrying Gas1* could form MVBs but, importantly, Gas1* is not internalized into ILVs. Since we did not observe GFP-Gas1*GPI on the vacuolar membrane in ESCRT mutants, a phenotype often observed for vacuolar substrates in this mutant background (Cabrera et al., 2013), we propose that ESCRT components on endosomes that carry specific misfolded proteins such as Gas1* could have additional functions besides catalyzing membrane invaginations for the generation of MVBs. In support of such a scenario, ESCRT components have been shown to affect endosome-vacuole/lysosome fusion in yeast and in mammals (Karim et al., 2018; Tumbarello et al., 2012).

**Figure 7.**
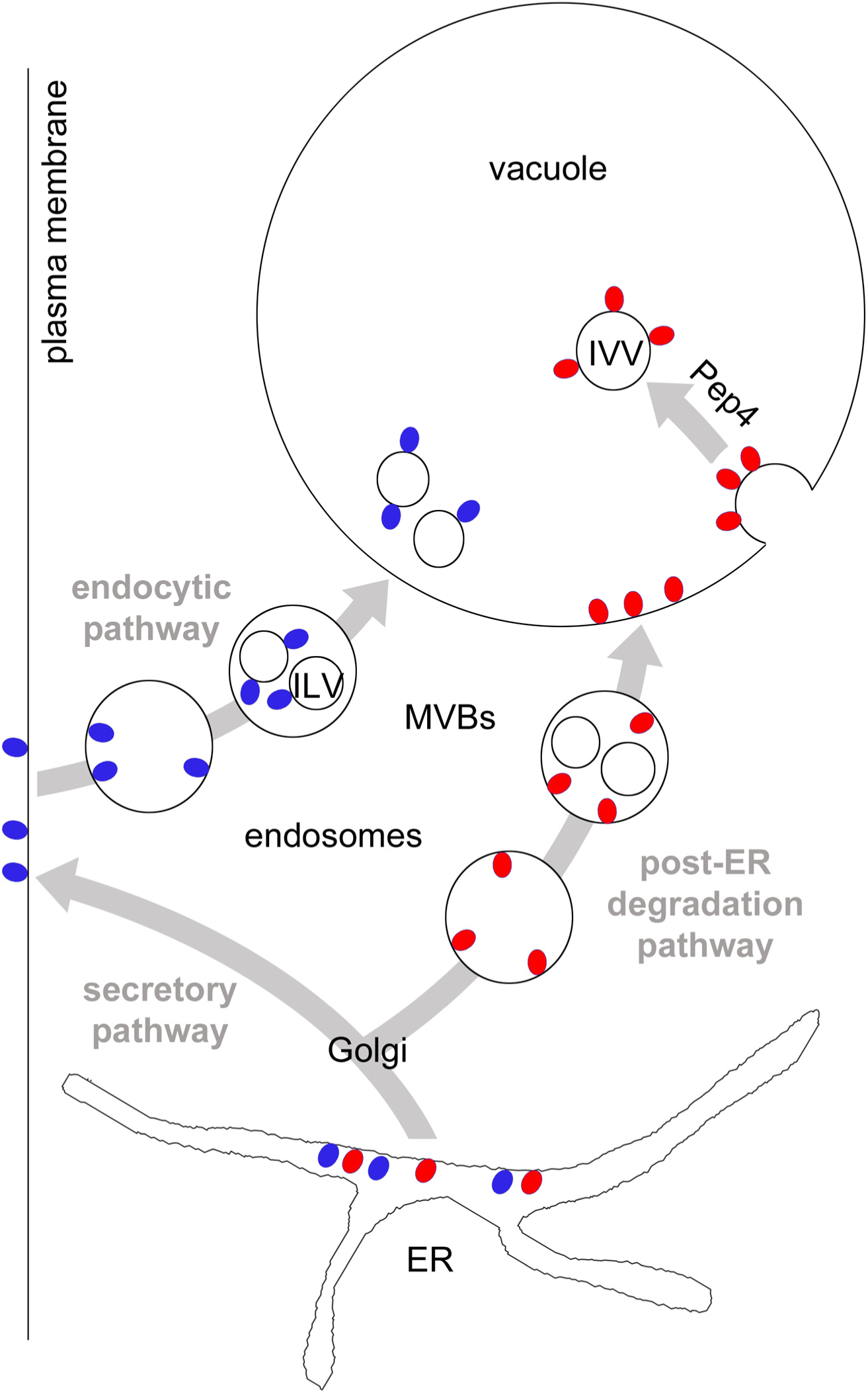
Model for cellular trafficking pathways of GPI-APs to the vacuole and the sites of protein internalization. Red and blue circles represent GPI-APs. They are synthesized at the ER and are exposed entirely to the luminal side. After ER export, misfolded GPI-APs (red circles) are diverted to endosomes, most likely from the Golgi. Lack of adaptors will prevent internalization into ILVs and after fusion of endosomes with the vacuole, GPI-APs will be exposed to the luminal side of the vacuolar membrane. Rapid internalization of GPI-APs from the vacuolar membrane is Pep4-dependent and leads to the generation of membrane-bound intravacuolar vesicles (IVV) which will be lysed in an Atg15-dependent manner. GPI-APs that are not detected by Golgi quality control mechanisms, including correctly folded species (blue circles), are routed to the plasma membrane, bypassing endosomes. Upon nutritional stimulus (MacDonald et al., 2015) or due to quality control mechanisms operating at the plasma membrane (Zavodszky and Hegde, 2019) they are endocytosed and routed to the vacuole/lysosome for degradation. Wild type Gas1 internalizes from late endosomes into ILVs, a process that is dependent on Cos proteins that operate in this pathway (MacDonald et al., 2015). See text for more details.

Second, we found that the modified reporter GFP-TMD-Gas1*GPI internalizes directly from the vacuolar membrane into Atg15-sensitive intravacuolar vesicles (IVVs); a process that bears some resemblance to microautophagy (Figure 7). Previously known cellular components with functions in various types of microautophagy in yeast, such as the VTC complex and the EGO complex, did not affect internalization of GFP-Gas1*GPI. Therefore we propose that the mechanism that we describe in this paper differs mechanistically in various ways from formerly known processes (Uttenweiler et al., 2007; Dubouloz et al., 2005). We observed accumulation in the vacuolar membrane with misfolded GPI-APs as well as with the reporter GFP-TMD-Gas1*TMD, which lacks a GPI anchor; thus, this phenomenon is not GPI-anchor-specific. Our original reporter GFP-Gas1*GPI is exposed exclusively to the luminal side of the organelles through which it traffics on its way to the vacuole and therefore can neither be ubiquitinated nor have direct access to the cytosolic ESCRT machinery. The observed absence of internalization of GFP-Gas1*GPI from endosomes suggests that the mechanism that is operational in the endocytic pathway and that relies on Cos proteins for ubiquitination of GPI-APs in *trans* is not accessible in the post-ER degradation pathway (Figure 7 and (MacDonald et al., 2015)). It could be possible that for substrates that are routed to the vacuole via the post-ER degradation pathway, only *cis* ubiquitination (i.e. direct ubiquitination of substrates) will lead to protein internalization by the ESCRT machinery. For instance, Wang et al. showed that the misfolded transmembrane protein Wsc1* is ubiquitinated on its cytosolic tail after diversion from the Golgi to endosomes and is subsequently internalized into ILVs before being delivered to the vacuole (Wang et al., 2011). In contrast, substrates that travel along the post-ER degradation pathway and cannot be *cis*-ubiquitinated, such as misfolded GPI-APs, would depend on the mechanism for internalization that we describe in this paper and that occurs from the vacuolar membrane (Figure 7). An extension of this model would suggest that endosomes that carry cargo from the plasma membrane and those that carry cargo from the post-ER degradation pathway might only have restricted or no mixing capacity (Figure 7). The small but reproducible fraction of GFP-Gas1*GPI that is still visible inside the vacuolar lumen in *Δpep4* cells could reflect the extent of endosomal mixing. Alternatively, the fraction of GFP-Gas1*GPI in the vacuolar lumen in *Δpep4* cells could reflect the small amount of the reporter that is first routed to the plasma membrane and then endocytosed, in which case it would encounter Cos proteins and internalize into ILVs.

In contrast to GFP-Gas1*GPI, the reporter GFP-TMD-Gas1*GPI does contain cytosolically exposed GFP with access to the ubiquitination machinery and the ESCRT-mediated internalization machinery from endosomes, yet we found it is not internalized from endosomes either. This is likely because GFP is not ubiquitinated *in vivo,* and its presence was furthermore shown to have inhibitory effects for the ubiquitination of neighboring domains (Chromotek, 2014; Baens et al., 2006).

Third, we discovered a previously unknown cellular mechanism for the regulation of vacuolar membrane dynamics. Our results show that the internalization of misfolded proteins can occur directly from the vacuolar membrane in dependence on the activity of the conserved vacuolar endoprotease Pep4 (related to Cathepsin D in mammals). This provides a novel general concept and suggests that a proteolytic event can be a molecular switch for the regulation of membrane invaginations. It is important to note that the deletion of Pep4 does not inhibit other processes that involve a dynamic rearrangement of the vacuolar membrane, such as the fusion of the vacuole with MVBs or the fusion with autophagosomes. This supports the conclusion that the defect in membrane dynamics in *Δpep4* cells is specific. We currently envision two major scenarios that could explain such an effect and that can be considered as testable hypothesis for future work. Both would depend on a Pep4-catalyzed proteolytic event inside the vacuole. In one scenario, Pep4 activity inside the vacuole creates a signal that would be transmitted across the vacuolar membrane for the modulation of cellular activities that regulate membrane invaginations from the cytosolic side, for instance through the ESCRT machinery. ESCRT components have recently been shown to have a role in inducing membrane invaginations from the vacuole, but the precise mechanisms are still unclear (Oku et al., 2017). Such a function of ESCRT components for the invagination of our reporters from the vacuolar membrane could be masked in our experimental approach, since all ESCRT mutants also affect the trafficking of the reporters to the vacuolar membrane and trap them at the PVC. A second and alternative scenario for how Pep4 could affect membrane invagination would be even more intriguing and could suggest that a Pep4-catalyzed proteolytic event inside the vacuolar lumen directly affects membrane bending from inside the vacuole. For instance, proteolysis of bulky luminal domains of vacuolar transmembrane proteins could facilitate the generation of the curvature at the neck of an inward budding vesicle. Either way, it will be interesting to unravel the underlying cellular mechanisms that link Pep4 activity to vacuolar membrane invaginations.

In summary, our data demonstrate the existence of a post-ER degradation pathway that is linked to a novel mechanism for protein internalization from the vacuolar membrane. Intriguingly, the activity of the conserved vacuolar/lysosomal protease Pep4 is required for this process and suggests that membrane dynamics can be regulated by proteolysis.

## Acknowledgements

We thank Antonio Miranda, Maya Schuldiner and Pedro Carvalho for critical reading of the manuscript and for many helpful comments. We are grateful to Maya Schuldiner for providing data concerning the intracellular localization of predicted yeast vacuolar proteases. This work was supported by grants of the Ministry of Economy and Competitiveness (grant BFU2016-78265-P) and the Erasmus+ program to Z.M. (HR ZAGREB01).

## Author contributions

L.L. and V.G. designed experiments, L.L., Z.M. and V.G. conducted experiments, L.L. and V.G. evaluated data, V.G. wrote the paper.

## Declaration of interest

The authors declare no competing interests.

## Material and Methods

### Yeast strains used in this study

A detailed list of yeast strains used in this study is found in the supplementary Table S1.

### Construction of plasmids used in this study

Unless stated otherwise, all constructs used in this study were expressed from centromeric or integrative plasmids under the control of the endogenous Gas1 promoter. Plasmid markers are indicated in the list of yeast strains which is appended as supplemental Table S1. The generation of constructs for the expression of GFP-Gas1*GPI (LLp14) and GFP-Gas1*TMD (LLp18) from integrative plasmids has been described previously (Sikorska et al., 2016). The construct for the expression of mRFP-Atg8 (LLp07) was a gift from Fulvio Reggiori. VGp115 is pESC-URA from Agilent Technologies. For the expression of GFP-Gas1*GPI from a centromeric plasmid, the coding sequence was released from LLp14 using XmaI and SacI and subcloned into pRS316 yielding LLp30. The construct expressing Gas1wtGFP-GPI from a centromeric plasmid (LLp66) is a gift from Laura Popolo. For the generation of the construct GFP-CPY*GPI (VGc409) the largest part of the coding sequence for Gas1* was replaced with the one for CPY*, maintaining only the N-terminal Gas1 signal sequence and the C-terminal Gas1 GPI anchor attachment site. First, the coding sequence for 81 amino acids downstream of the internal restriction site BsrGI in a construct expressing Gas1* was removed using the QuikChange protocol (Agilent), yielding VGc389. Subsequently, the coding sequence of CPY* was amplified from pDN431 (Ng et al., 2000) using the primers 5’GCGCATATGTCATTGCAAAGACCGTTG and 5’GCGTGTACATAAGGAGAAACCACCGTG. The PCR product was cut with NdeI and BsrGI and was used to replace the Gas1* fragment released from VGc389 after cutting with the same enzymes, yielding VGc392. To insert the coding sequence for GFP, the GFP sequence was amplified from pKT128 (EUROSCARF collection) with the primers 5’GCGACGCGTTCTAAAGGTGAAGAATTATTC and 5’GCGACGCGTTTTGTACAATTCATCCATACC. The PCR product was cut with MluI and inserted into the internal MluI site in the remaining Gas1 coding sequence, just downstream of the N-terminal signal sequence, yielding VGc395. Finally, the construct was digested with XmaI and SacI and the put into pRS306 for expression from an integrative plasmid, yielding VGc409. The construct GFP-TMD-Gas1*GPI (VGp204) was expressed from a centromeric plasmid under the control of the TDH3 promoter. An initial fusion construct coding for GFP-TMD-Gas1* with the TMD from the yeast ER protein Ted1 that is predicted to orient with the desired topology in the ER membrane (Ncyt) and several internal and adjacent restrictions sites was synthesized by an external company. The TDH3 promoter was amplified from the yeast strain BY4741 using the primers 5’GCGCCCGGGCAGTTCGAGTTTATCATTATC and 5’CGCGCTAGCCATTTTGTTTGTTTATGTGTGTTT and put in front of the coding sequence using the restriction enzymes XmaI and NheI. Finally, the entire construct together with the TDH3 promoter was subcloned into the centromeric yeast expression vector pRS316 using XmaI and SpeI, yielding VGp204. The construct was subcloned into into the centromeric yeast expression vector pRS315 using XhoI and SacI, yielding VGp338. The construct GFP-TMD-Gas1*TMD (VGp206) was generated by using a modified version of the QuikChange protocol (Agilent). The construct VGp204 was used as template and the single primer 5’CAGCTTCATCTTCATCTTCTTCGCGAAAGCAAGCTGCCACCAACGTTAAAGC was used to introduce the point mutation N528Q which destroys the GPI anchor attachment site. The construct mRFP-Ufe1 (VGc344) was expressed from a centromeric plasmid under the control of the TDH3 promoter. The generation of a construct expressing Ufe1 under the control of the TDH3 promoter (VGp172) has been described previously (Lemus et al., 2016). To insert mRFP in the N-terminal region of Ufe1, the coding sequence of mRFP was amplified by PCR from a plasmid containing mRFP-Atg8 (Mari et al., 2010) using the primers 5’CGCGCTAGCATGGCCTCCTCCGAGGACG and 5’CGCGCTAGCGGCGCCGGTGGAGTGG. The PCR product was cleaved with NheI and inserted into VGp172 cut with XbaI, yielding VGc344. To express myc-tagged Pep4 under the control of its endogenous promoter, the ORF of Pep4 including the 300bp upstream region was amplified from BY4741 using the primers 5’GCGACTAGTGGATTTATATAAAAAGCCATACTTC and 5’CGCGTCGACAATTGCTTTGGCCAAACCAAC. The PCR product was cut with SpeI and SalI and ligated into the expression vector pESC-URA (Agilent) cut with the same enzymes, yielding VGc448. The catalytically inactive Pep4 mutant was generated by using a modified version of the QuikChange protocol (Agilent). The construct VGc448 was used as template and the single primer 5’AAAACTTCAAGGTTATTTTGAAGACTGGATCCTCAAACCTTTGGGTTCCAAG was used to introduce the point mutation D109K and a diagnostic restriction site, yielding VGc451.

### GFP-processing assays

Cells were grown overnight, diluted to OD 0.2 and regrown for 5 hours. Removed aliquots were lysed by alkaline treatment (Kushnirov, 2000), resuspended in a cell-density normalized volume of loading buffer, followed by SDS-PAGE and Western Blotting using anti-GFP antibody (Roche), HRP-conjugated anti-mouse secondary antibody (Roche) and ECL (PIERCE) as substrate. Images were taken with a LAS-3000 mini imaging system (Fujifilm) and bands were quantified using Multi-Gauge software (Fujifilm).

### Staining with FM4-64

To visualize vacuolar membranes, cells were stained with FM4-64, according to (Vida and Emr, 1995). Briefly, log-phase cells were incubated with 8μM FM4-64 (8mM stock in DMSO) for 30 min at 30 °C in the dark. Cells were washed and incubated for additional 90 min prior to live cell fluorescence microscopy using the RFP filter.

### Fluorescence microscopy

Exponentially growing cells were washed with PBS and immediately analyzed by fluorescence microcopy. Cells were observed with a LEICA DMi8 microscope equipped with a 100x/1.4 oil Plan-Apo immersion lens and a DIC prism and polarizer for Normarksi imaging. Images were acquired using a Hamamatsu C1 3440-20CV camera and the LASX controller software (Leica).

### Confocal live cell microscopy

Exponentially growing cells were washed, resuspended in PBS and scanned with a Laser Scanning Confocal Microscope from Zeiss (LSM 7 Duo) equipped with a BiG (binary GaAsP) module using a Plan-Apochromat objective 63x/1,40 Oil DIC. A 488nm Argon Laser (GFP) and a 561nm Helium-Neon Laser were used (mRFP).

---

**Figure S1:**
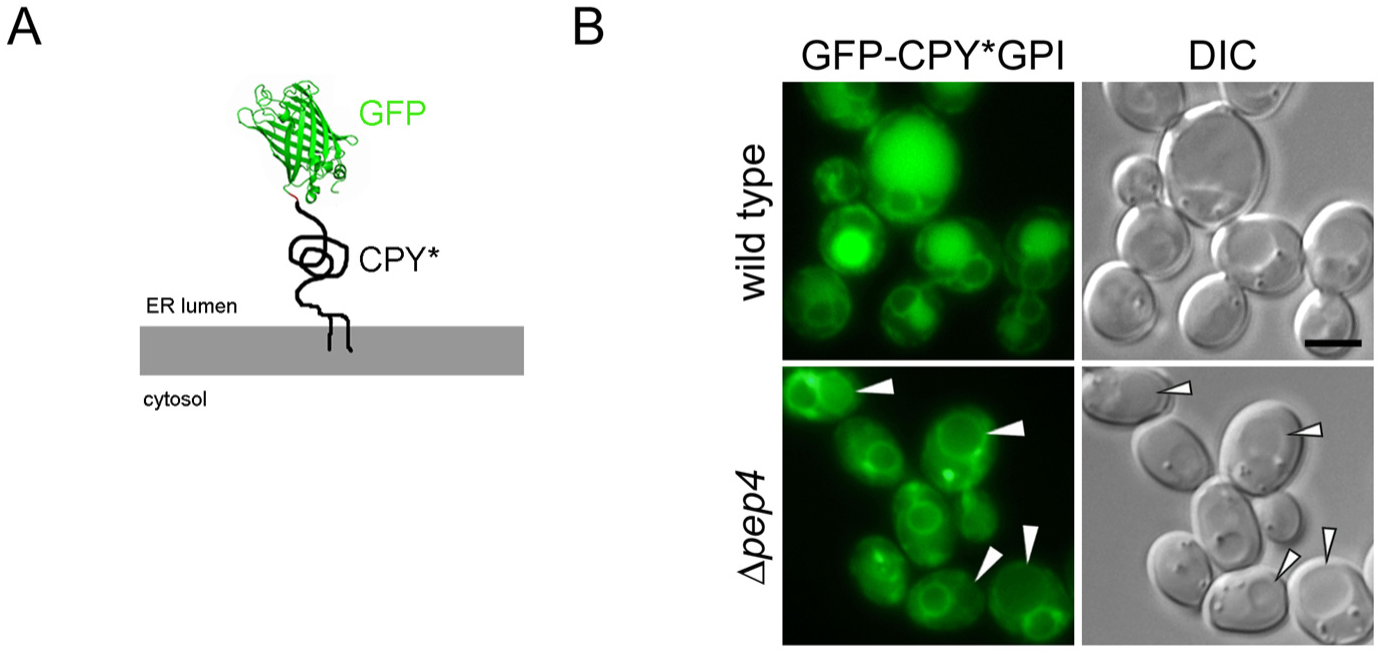
The misfolded GPI-anchored protein GFP-CPY*GPI is routed to the limiting membrane of the vacuole prior to degradation. In order to test whether an additional misfolded GPI-AP is routed to the limiting membrane of the vacuole prior to degradation we generated GFP-CPY*GPI. In this construct, CPY*, a well-studied misfolded soluble ER-associated protein degradation (ERAD) substrate, was fused with GFP at its N-terminus, downstream of its signal sequence, and with the 57 C-terminal amino acids of Gas1 that contain the GPI anchor attachment site (Sikorska et al., 2016). **(A)** Schematic representation of the fusion construct GFP-CPY*GPI. The C-terminal GPI anchor extends into the luminal leaflet of the ER membrane. **(B)** Wild type cells and *Δpep4* cells expressing genomically integrated GFP-CPY*GPI under the control of the GAS1 promoter were analyzed by live cell fluorescence microscopy and DIC microscopy. Scale bar: 5μm. When expressed in wild type cells, staining of the perinuclear ER and the vacuole was visible. Cells with Pep4 deleted showed strongly reduced staining of the vacuolar lumen and showed mainly staining of the limiting membrane of the vacuole, marked with arrow heads. Related to Figure 1.

**Figure S2:**
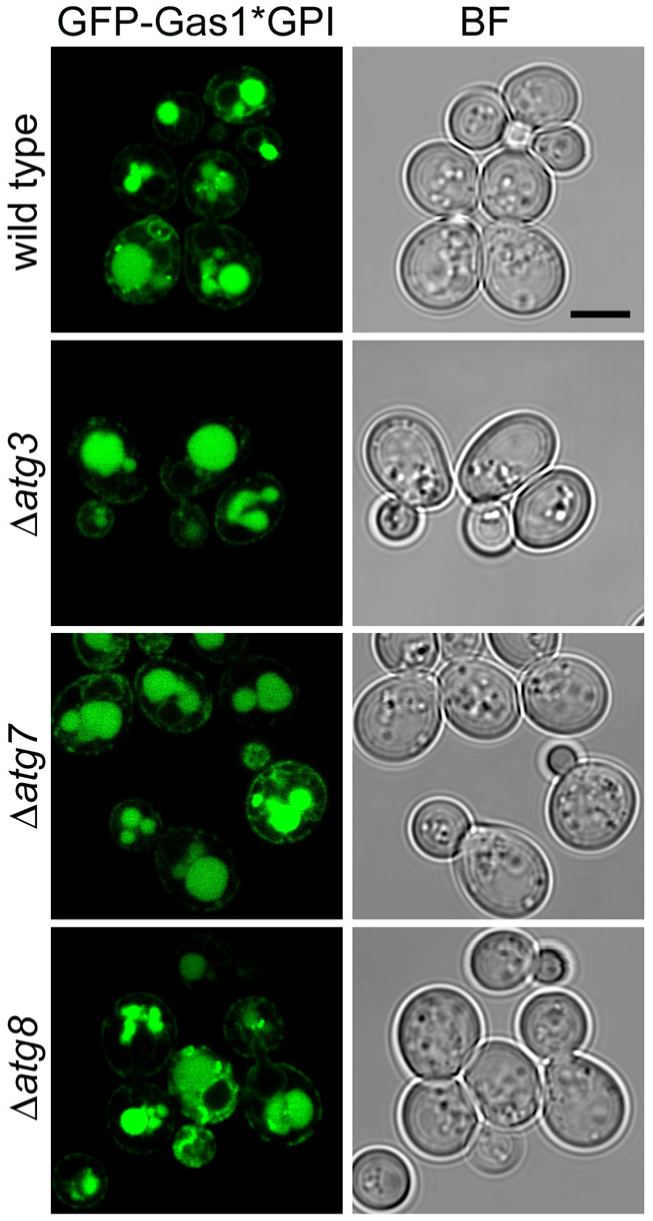
Routing of GFP-Gas1*GPI to the vacuole is unaffected in autophagy mutants. Wild type cells and the indicated deletion mutant cells expressing GFP-Gas1*GPI from a CEN plasmid under the control of the GAS1 promoter were analyzed by live cell laser scanning confocal fluorescence microscopy. Scale bar: 5μm. BF = bright field. A large fraction of GFP-Gas1*GPI is found inside the vacuole in wild type cells as well as in cells that are defective in generating autophagosomes, showing that autophagy is not involved in routing of the reporter to the vacuole. Related to Figure 2.

**Figure S3:**
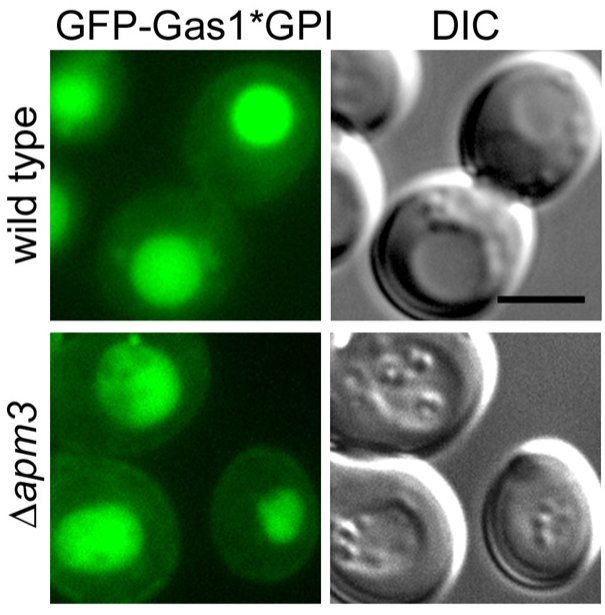
Routing of GFP-Gas1*GPI to the vacuole is unaffected in Δ*apm3* cells. Wild type cells and *Δapm3* cells expressing GFP-Gas1*GPI from a CEN plasmid under the control of the GAS1 promoter were analyzed by live cell fluorescence microscopy in combination with DIC microscopy. Scale bar: 5μm. GFP-Gas1*GPI localization in *Δapm3* cells at steady state is indistinguishable from that in wild type cells, showing that the AP-3 pathway is not involved in routing of the reporter to the vacuole. Related to Figure 3.

**Figure S4:**
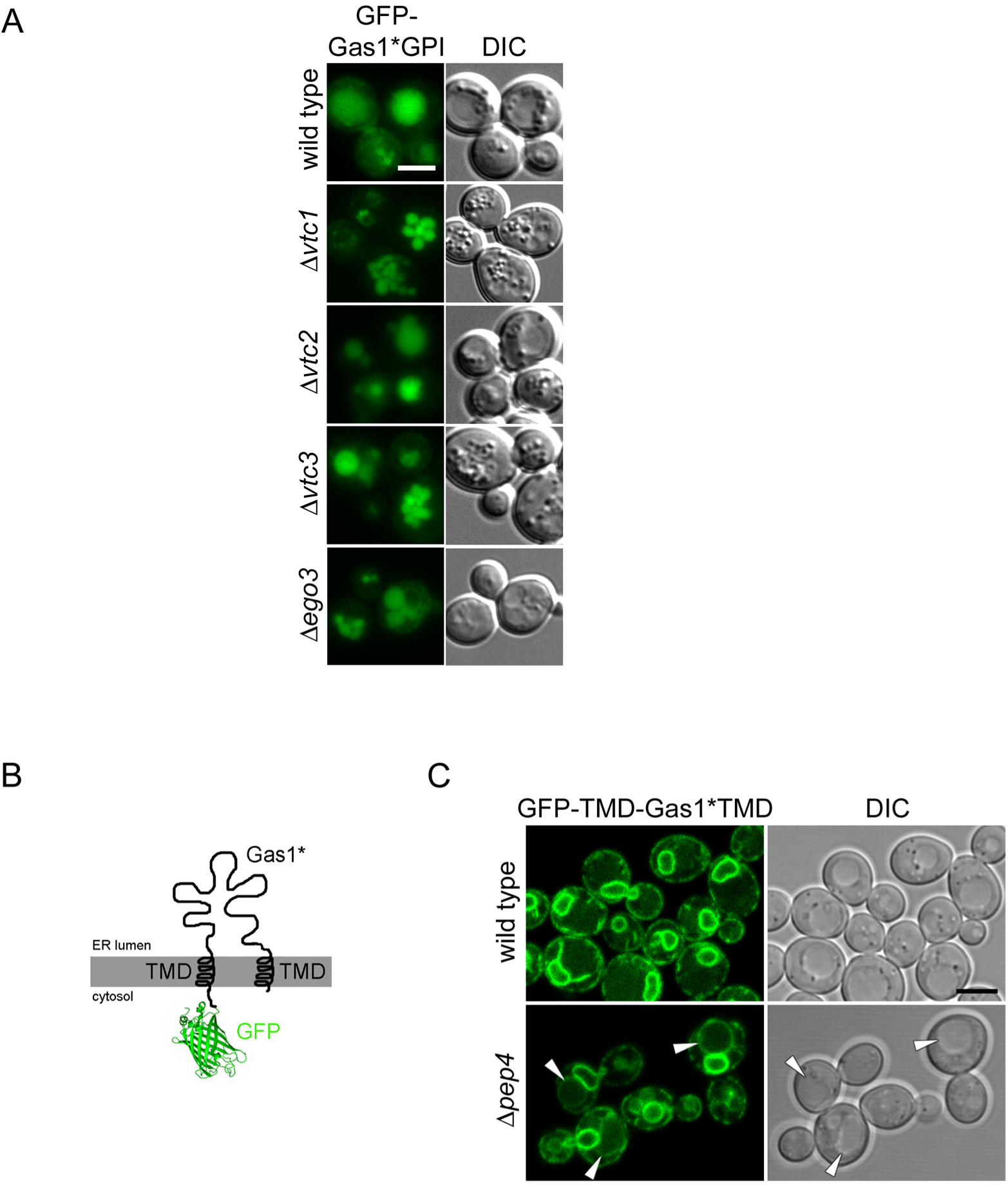
Live cell fluorescence microscopy linked to reporter internalization from the vacuolar membrane. **(A)** Vacuolar uptake of GFP-Gas1*GPI is unaffected in mutants previously shown to be involved in microautophagy. Wild type cells and the indicated mutants expressing GFP-Gas1*GPI from a CEN plasmid under the control of the GAS1 promoter were analyzed by live cell fluorescence microscopy and DIC microscopy. Scale bar: 5μm. None of the mutants previously known for their roles in microautophagy (Uttenweiler et al., 2007; Dubouloz et al., 2005) showed an enrichment of GFP-Gas1*GPI in the vacuolar membrane, demonstrating that they are not involved in internalization of the reporter into the vacuole**. (B)** Schematic representation of the construct GFP-TMD-Gas1*TMD. The construct is identical to GFP-TMD-Gas1*GPI (Figure 4A) with the exception that the C-terminal GPI anchoring signal has been destroyed with a single point mutation, preventing the exchange of the C-terminal TMD for a GPI anchor. **(C)** GFP-TMD-Gas1*TMD is efficiently retained inside the ER. Wild type cells and*Δpep4* cells expressing GFP-TMD-Gas1*TMD from a CEN plasmid under the control of the GAS1 promoter were analyzed by live cell laser scanning confocal fluorescence microscopy and DIC microscopy. Scale bar: 5μm. Vacuolar staining is weak in wild type cells, and the perinuclear ER is very prominently labeled. These data support the idea that the presence of the C-terminal GPI anchor is responsible for the routing of GFP-TMD-Gas1*GPI (Figure 4A) to the vacuole, as expected. Importantly, the minor fraction of GFP-TMD-Gas1*TMD that is still routed to the vacuole is seen enriched at the vacuolar membrane in *Δpep4* cells (marked with arrow heads). Related to Figure 4.

**Figure S5:**
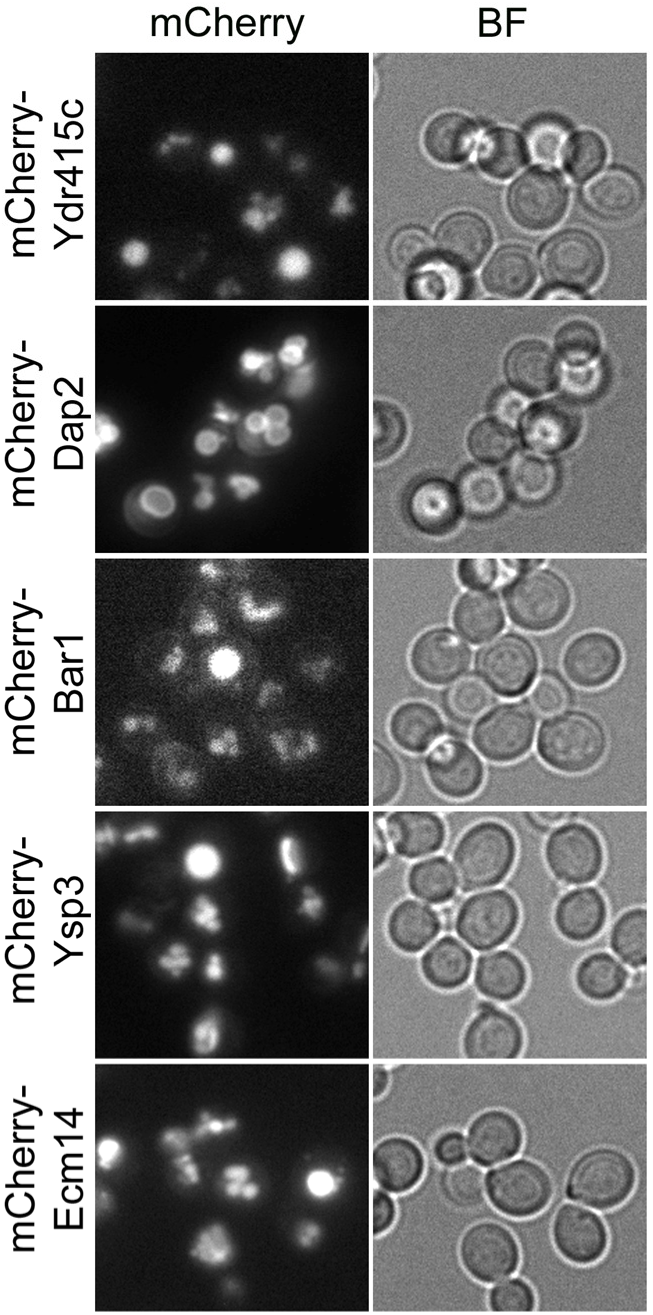
Vacuolar localization of predicted proteases. The indicated fusion proteins are expressed in wild type cells from the genomic locus under the control of the TEF2 promoter and contain an N-terminal Kar2 signal sequence followed by mCherry and the protein of interest. Images are a courtesy of the Schuldiner lab (Weizmann Institute, Rehovot, Isreal) and have been acquired in the course of a previously published high through-put study using an automated microscopy setup (ScanR system, Olympus) as described (Yofe et al., 2016). Fusion proteins of Ydr415c, Bar1, Ysp3 and Ecm14 localize to the vacuolar lumen at steady state, whereas Dap2 is targeted to the vacuolar membrane. Related to Figure 5.

## Legends for supplemental movies

**Movie S1:** *Δatg15* cells expressing GFP-TMD-Gas1*GPI show GFP-containing intravacuolar mobile structures. The movie covers 50 seconds with one frame taken every 2 seconds. Related to Figure 4.

**Movie S2:** *Δpep4Δatg15* cells expressing GFP-TMD-Gas1*GPI lack the appearance of GFP-containing intravacuolar mobile structures. The movie covers 50 seconds with one frame taken every 2 seconds. Related to Figure 4.

**Table S1.**
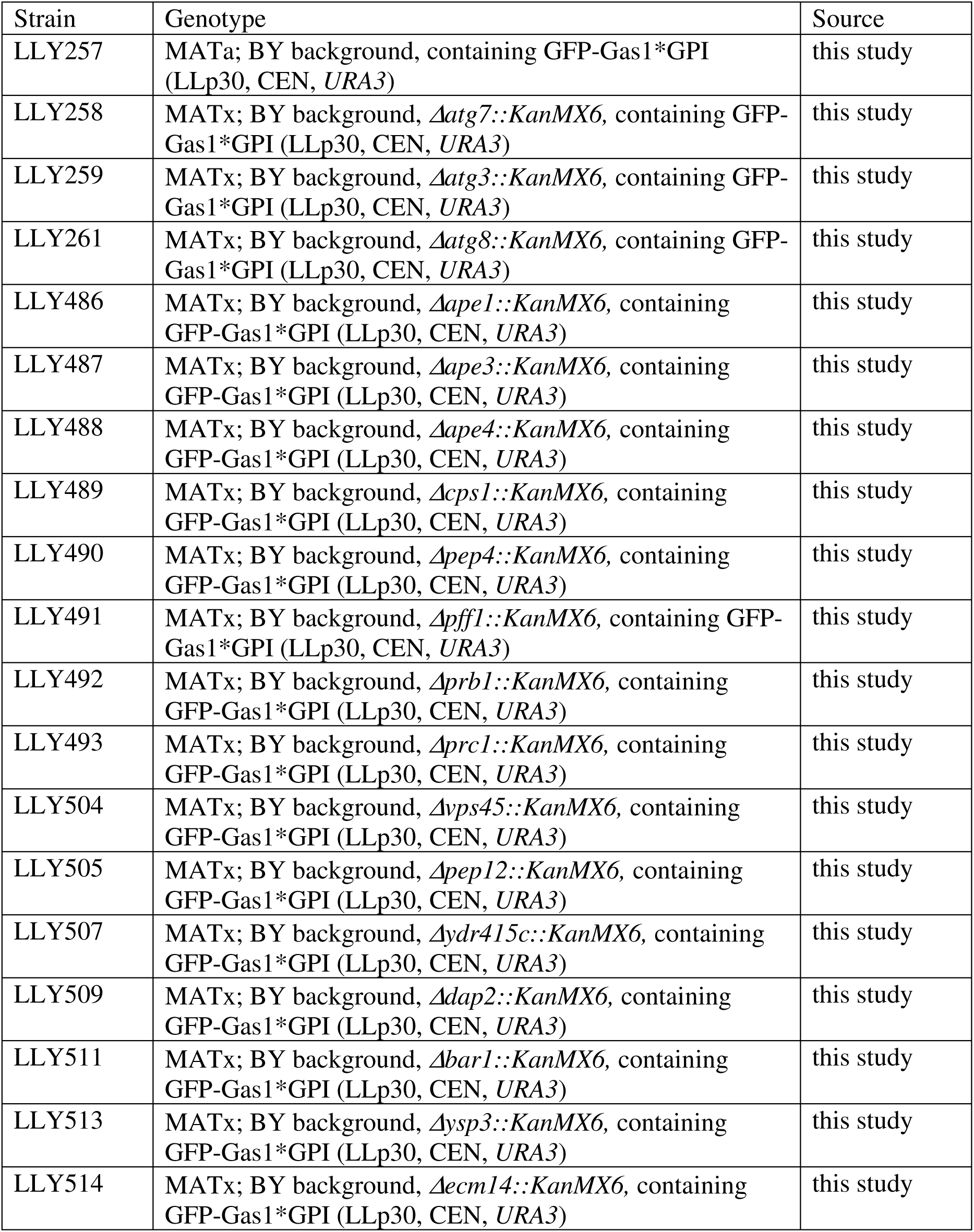
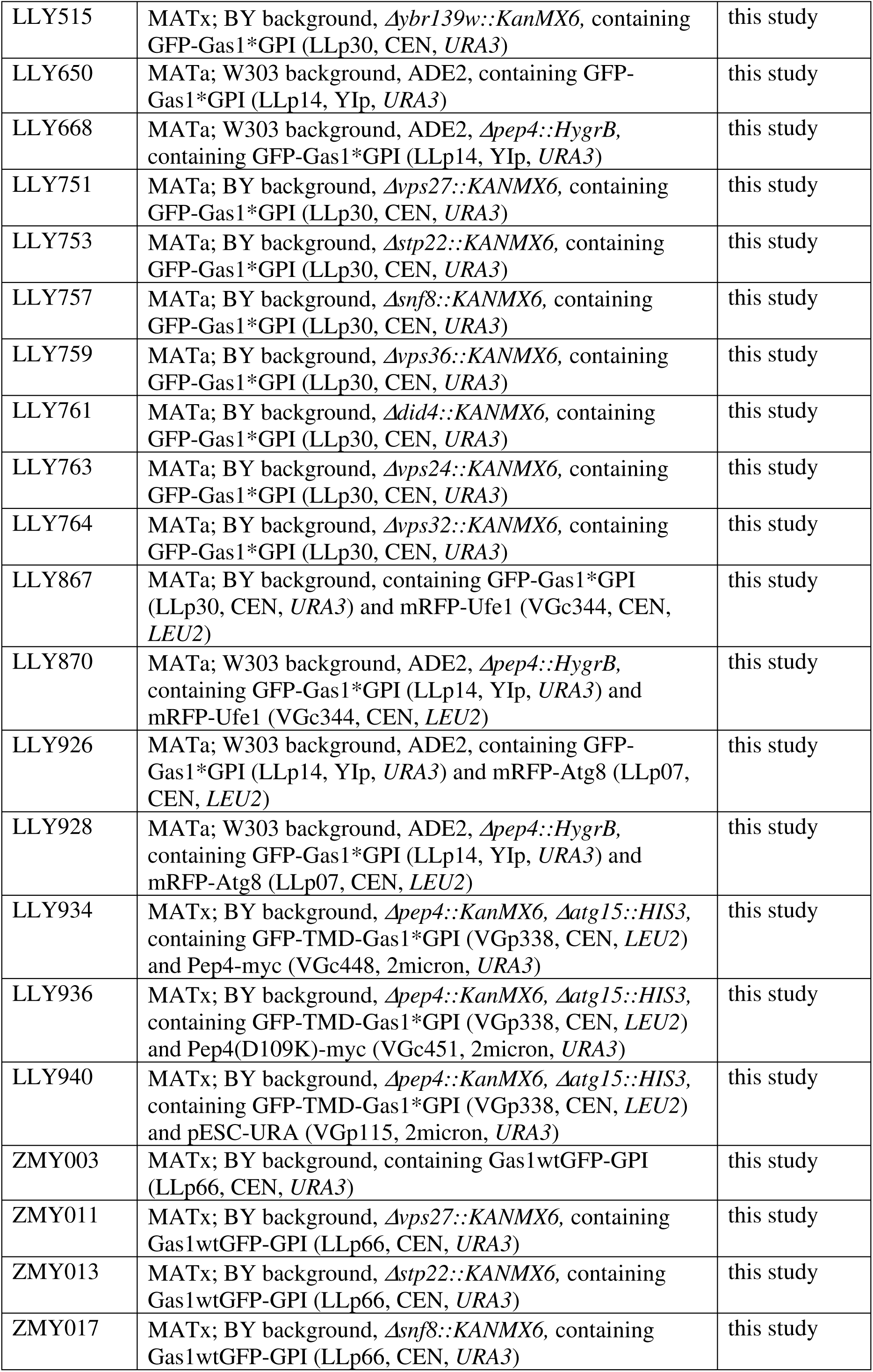
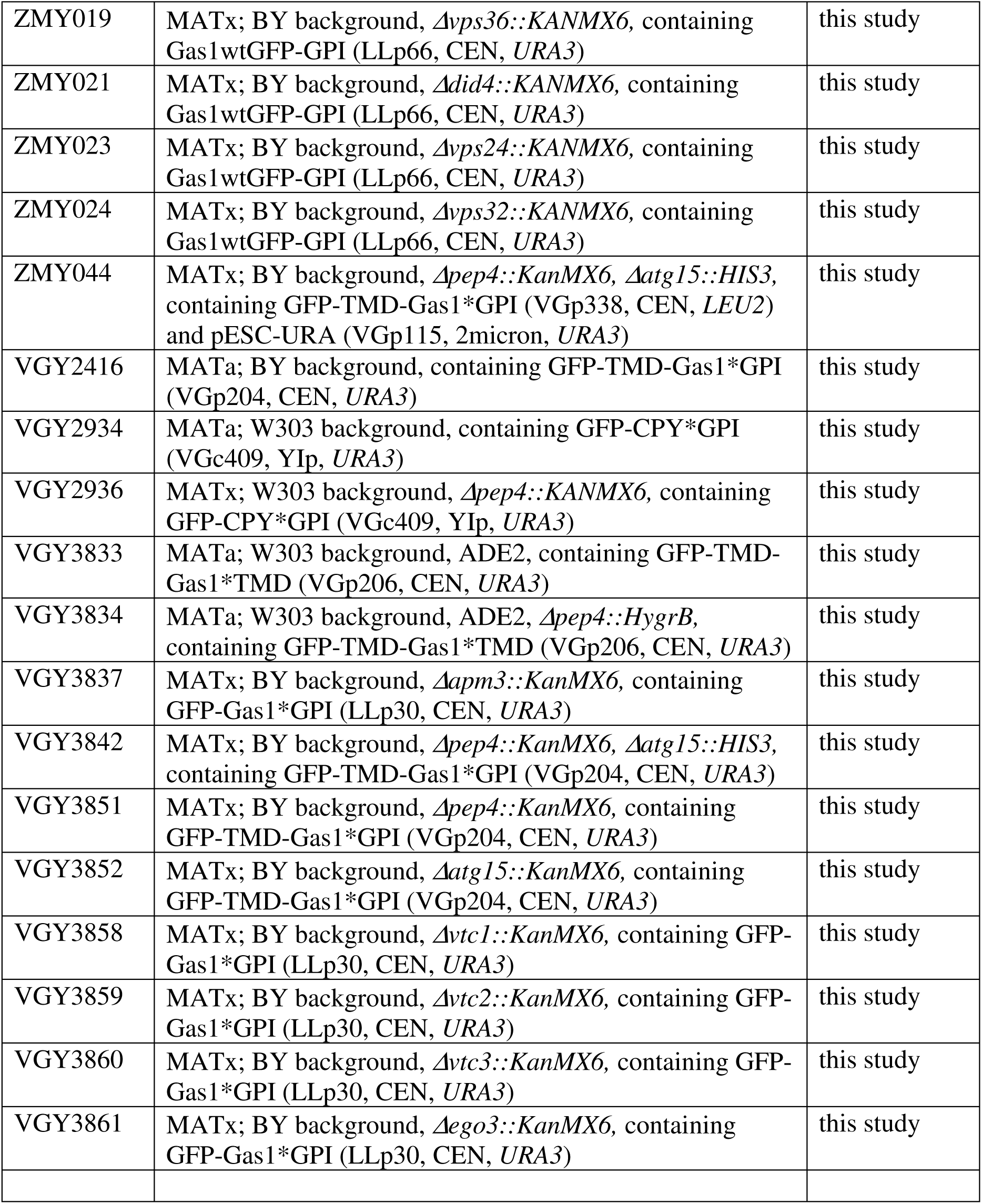
Yeast Strains used in this study. Chromosomal tagging of proteins, gene deletions and integrations were performed (if needed) using PCR in combination with standard homologous recombination techniques.

